# Actin filament assembly driven by distributive polymerases clustered on membrane surfaces

**DOI:** 10.1101/2024.11.26.625540

**Authors:** R. Dyche Mullins, Jane Kondev, Kristen Skruber

## Abstract

Actin filaments created by the Arp2/3 complex form branched networks, that grow and push against cellular membranes. We employ theory and simulation to describe how membrane surfaces accelerate filament assembly via clustering of proteins that bind actin monomers and/or profilin-actin complexes. Briefly, thermal fluctuations drive filament tips on constrained, two-dimensional random walks across the membrane, where they encounter multiple actin-charged polymerases. At low actin concentrations, filament elongation is limited by delivery of monomers to the membrane surface; at high actin concentrations, elongation depends on how quickly fluctuating filaments search the membrane. Using experimentally measured parameter values we conclude that surface-mediated polymerization can outpace solution-mediated elongation, even at high actin concentrations (>200 µM). The finite time required for profilin dissociation decreases the advantage conferred by surface-associated polymerases, but only in the absence of force. Load forces enhance the effect of surface polymerases, which can both accelerate elongation and increase the force required to stall filament assembly.

## INTRODUCTION

In eukaryotic cells the actin cytoskeleton moves, shapes, and reinforces cellular membranes. In fact, most actin filaments are created at membrane surfaces (Welch, 2004) where their rapid assembly can generate pushing forces (Mogilner, 1996) that support many cellular processes. Branched actin networks created by the Arp2/3 complex, for example, harness the growth of multiple actin filaments to produce large-scale compressive forces that drive pseudopod protrusion (Bisi, 2013), endocytosis (Mooren, 2012), phagocytosis (Insall, 2009), cell-cell adhesion (Efimova, 2018), cell fusion (Richardson, 2007), healing of membrane ruptures (Clark, 2009), and assembly of autophagosomes (Kast, 2015). Despite the biological importance of actin filament assembly in branched networks, we lack realistic, quantitative models to describe how this process works in living cells or complex, reconstituted systems *in vitro*.

A common assumption is that elongation of cellular actin filaments mirrors that of purified actin *in vitro* (Pollard, 2000). Specifically, purified actin forms polarized filaments that elongate when soluble monomers bind to one of their ends. The rates of elongation differ at the two filament ends and also differ depending on whether incoming monomers are bound to ATP or ADP (Pollard, 1986; Fujiwara, 2007). In the cytoplasm most monomeric actin is likely bound to ATP (Rosenblatt, 1995) and complexed with the monomer-binding protein, profilin (Kaiser, 1999). When a profilin-actin complex binds to a free barbed end, the newly attached actin monomer undergoes a rapid conformational change that accelerates profilin dissociation (Pollard, 1984). At relatively low concentrations, therefore, profilin has little effect on the rate of elongation at the fast-growing (*barbed*) end of the filament, but strongly inhibits elongation from the slow-growing (*pointed*) end (Tilney, 1983; Funk, 2019). Based on these observations, the major mode of actin filament elongation in cells is commonly assumed to be the rapid, diffusion-limited (Drenckhahn, 1986) binding of soluble ATP-actin monomers and/or profilin-ATP-actin complexes to free barbed ends (Pollard, 2000).

Many quantitative models of cellular actin assembly explicitly assume that filaments elongate from soluble building blocks, according to rate constants measured *in vitro*. (Schaus, 2007; Garner, 2022). Unfortunately, few spatially correlated measurements of soluble actin concentrations and filament elongation rates have been made in living cells, and the available data suggest that the relationship between monomer concentration and filament elongation is not so simple (e.g. Koestler, 2009). At least three additional factors affect the rate of filament elongation in cells: (i) physical forces, (ii) the activity of actin polymerases, and (iii) the dissociation of profilin from barbed ends. Physical forces acting on the end of a filament can dramatically slow its growth (Hill, 1982; Mogilner, 1996; Li, 2022); while the activity of actin polymerases can accelerate elongation by several-fold (Romero, 2004; Kovar, 2006; Breitsprecher, 2008; Hansen, 2010; Bieling, 2018). And, ultimately, the rate of profilin dissociation sets an upper limit on the rate of filament elongation (Funk, 2019).

Models of formin-generated cytoskeletal structures often explicitly incorporate the polymerase activity of formin-family proteins, especially that of the most highly processive formins (Mohapatra, 2015). In contrast, the effect of force is sometimes incorporated into quantitative models of branched actin network assembly (e.g. Schaus, 2007), but the influences of polymerase activity and profilin dissociation are almost always neglected. This is a problem because the barbed ends of actin filaments in an Arp2/3-generated network generally elongate against membranes coated with a high density of nucleation promoting factors of the WAVE/WASP family. In addition to activating the nucleation activity of the Arp2/3 complex, WAVE/WASP proteins also contain binding sites for monomeric actin and profilin-actin complexes that, when clustered on a surface, work together to create a distributive actin polymerase that can accelerate elongation of nearby filaments by several fold (Bieling, 2018).

Here we provide the mathematical framework for a more realistic description of how actin filaments grow in contact with polymerase-coated membranes. We calculate the rate of membrane-driven filament elongation as a function of both the soluble profilin-actin concentration and polymerase surface density. Using this result we then compute the relative contributions of soluble and membrane-associated actin monomers to overall filament elongation. Using measured parameter values and conservative assumptions, we calculate that more than 50% of actin subunits enter a branched network from surface-associated polymerases, even at high soluble actin concentrations (>200 µM). At cellular profilin-actin concentrations (∼100 µM) we estimate that ∼75% of actin is delivered to the network via the membrane surface. Filaments begin to interfere with each other and slow the rate of surface-mediated elongation when they come into close enough proximity for their membrane-interaction regions to overlap. Capping proteins, therefore, promote growth of branched actin networks (Hug, 1995; Mejillano, 2004; Iwasa, 2007; Akin, 2008) by decreasing inter-filament competition for surface-bound monomers. Interestingly, profilin dissociation decreases the growth advantage conferred by surface-associated polymerases, especially at high soluble profilin-actin concentrations. When profilin-limited filaments experience a load force, however, polymerase-coated surfaces again confer an advantage, one that increases with increasing force. Finally, under physiologically relevant conditions, we calculate that polymerase-coated membrane surfaces can increase the force required to stall the growth of a nearby actin filament by ∼1 pN.

## Results

In this section we develop a nanometer-scale model of an actin filament growing near a membrane coated with WAVE/WASP-family proteins. We describe the relevant properties of the membrane surface and the tip of a growing actin filament, and then use these properties to calculate the rate of surface-mediated filament elongation.

Three nucleation promoting factors —WAVE, WASP, and WASH— are conserved across eukaryotic phyla, and were likely inherited from the last common ancestor of all eukaryotes (Fritz Laylin, 2017). These proteins and their relatives cluster on membrane surfaces (Figure 1A, 1B) where they direct formation of branched actin networks by locally stimulating Arp2/3-dependent filament nucleation and by accelerating filament growth (Machesky, 1999; Bieling, 2018). The actin polymerase activity of WAVE/WASP-family proteins arises from two adjacent sequence motifs, a proline-rich region that binds multiple profilin-actin complexes and at least one WASP homology 2 (WH2) sequence that binds monomeric actin (Figure 1A). We characterize WAVE/WASP proteins on the membrane by specifying three key properties: (i) mode and stoichiometry of actin monomer binding; (ii) mobility on the membrane surface; and (iii) surface density.

**Figure 1.**
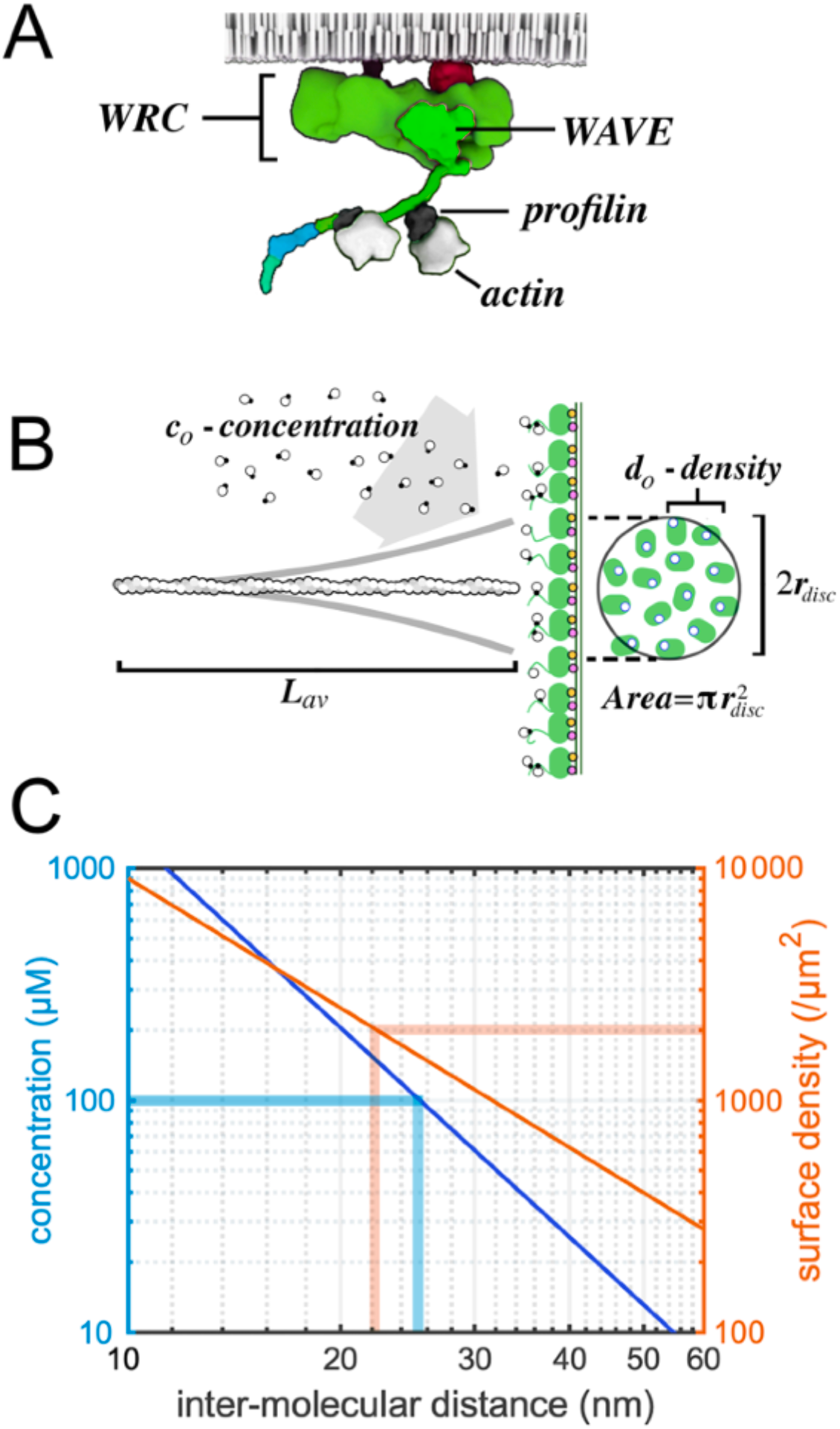
WAVE-family actin polymerases. (A) The WAVE Regulatory Complex (WRC) bound to two profilin actin complexes and attached to a membrane by small G-proteins. (B) Actin filament (length, *Lav*) growing against a WRC-coated membrane with binding sites for profilin-actin (black and white circles; density, *d*_*o*_,). Thermal fluctuation of the filament (thick gray lines) defines a membrane disc with (radius, *r*_*disc*_) containing multiple polymerases. (C) Log-log plot of solution concentration (blue line and left y-axis) and surface density (orange line and right y-axis) versus average inter-molecular spacing (x-axis). Thick blue lines mark 100 µM and 25 nm. Thick orange lines mark 2000/ µm^2^ and 22 nm.

### WAVE/WASP-family proteins on the membrane

Most monomeric actin in the cytoplasm is likely bound to profilin (Kaiser, 1999), so we will ignore the contribution to polymerase activity of the actin monomer-binding WH2 domains and consider only the profilin-binding, proline-rich regions. Human WAVE1 contains up to six profilin-actin binding sites with a range of measured affinities, from 2 µM to ∼200 µM (Bieling, 2018). It is unclear whether six profilins could ever bind WAVE at the same time but, *in vitro*, profilin:WAVE stoichiometries of at least 2:1 have been detected (Bieling, 2018). This result suggests that the density of polymerizable actin on a WAVE1-coated membrane is at least twice that of WAVE1 itself (Figure 1A).

WAVE proteins localize to membrane surfaces via interaction with small, rho-family G-proteins (Figure 1A)(Koronakis, 2011). In this analysis we ignore lateral motion of membrane-associated WAVE molecules for two reasons. Firstly, the measured diffusion coefficient of WAVEs on the plasma membrane of a motile cell is an order of magnitude lower than that of nearby actin filament tips (Millius, 2012; Mogilner, 1996). Secondly, our previous *in vitro* experiments demonstrated that lateral mobility of WAVE is not required for polymerase activity (Bieling, 2018).

Estimates of WAVE/WASP protein density on cellular membranes range from >1000/µm^2^ for WAVE at the leading edge of crawling neutrophils (Weiner, 2007) to 25,000/µm^2^ for WASP at sites of endocytosis in budding yeast (Arasada, 2011). We focus on the lower end of this range and the specific case of mammalian WAVE proteins. WAVEs function as part of a 5-subunit complex, whose overall shape can be approximated as a ∼19×8 nm prolate ellipsoid (Figure 1A; Chen, 2010), with a maximum packing density of ∼6,600/µm^2^ on a membrane surface (Figure 1B). Our analysis, therefore, concentrates on WAVE surface densities of 1,000-6,000/µm^2^. As a useful comparison (Huang, 2024), a surface density of 2000/µm^2^ reflects an average distance of ∼22 nm between WAVE molecules —the same inter-molecular spacing as a three-dimensional solution with a concentration of ∼150 µM (Figure 1C).

### Filament tip dynamics define a ‘footprint’ on the membrane

Assuming that WAVE molecules are relatively immobile, the region of membrane that contributes actin monomers to a nearby filament depends mainly on lateral mobility of the filament tip. From theoretical and experimental studies we know that thermal forces cause bending fluctuations in filaments growing against a WAVE-coated surface (Mogilner, 1996; Bieling, 2022). These bending fluctuations cause the tip of the filament to trace a constrained random walk across the surface of the membrane. The shape of the region defined by this random walk depends on the orientation of the filament relative to the membrane and the compressive force acting on it. The tip of an unloaded filament oriented normal to the membrane surface, for example, will trace out a circular disc (Figure 1B). This disc will contain a number of polymerase-bound actin molecules (*n*_*pol*_) that depends on its surface area and the local density of WAVE molecules. Compressive loading, however, will prevent the filament from straightening and, thus, decrease the frequency with which its tip visits the center of the disc. Under these conditions the envelope of tip positions becomes more annular. In any other orientation the filament tip will trace more elliptical and/or ‘croissant’ shapes. For simplicity, we begin by considering an unloaded filament oriented normal to the membrane surface. In this case the radius of the membrane interaction disc (*r*_*disc*_) depends only the length and stiffness of the filament.

Estimates of the average filament length in a functional branched actin network range from ∼60-300 subunits (∼150-800 nm; Svitkina, 1999; Akin, 2008; Koestler, 2009). Here, we assume an average filament length (*L*_*av*_) of ∼110 subunits (∼300 nm) from free barbed end to network-anchored pointed end. At this length the root-mean square (RMS) displacement of the filament tip due to thermal fluctuations will be ∼45 nm (Appendix B). If these filaments grow in proximity to a polymerase-coated membrane (Figure 1B) we define this RMS displacement as the radius of the membrane interaction disc (*r*_*disc*_) —i.e. the ‘footprint’ of the filament tip on the membrane surface. At densities of 1000-6000/µm^2^ the average number of WAVE molecules in this disc lies between 6 and 36. We limit our analysis to conditions under which the soluble profilin-actin concentration (*c*_*o*_) significantly exceeds the equilibrium dissociation constant of the two main profilin-actin binding sites on WAVE (2 µM and 4 µM; Bieling, 2018). In which case the total profilin-actin binding capacity of the membrane interaction disc (*n*_*pol*_) ranges from 12 to 72.

### Filament elongation

To calculate the rate of surface-mediated actin filament elongation we determine the rate at which thermal fluctuation of filament tips causes them to encounter membrane-associated actin monomers. This type of calculation produces a collision rate, analogous to the “Smoluchowski limit” on the rate of diffusion-limited molecular interaction. This approach ignores steric and electrostatic effects on elongation, which we discuss in a later section. We perform this computation in three different ways. First, we derive an analytical solution assuming that the filament tip undergoes a discrete random walk on a rectangular grid bounded by a circular perimeter (Appendix C). Second, we take a continuum approach similar to that of von Smoluchowski (1917). In this approach we calculate the diffusion-limited rate of monomer-filament interaction in two dimensions (Appendix D) and then determine the effect of imposing a circular boundary on tip diffusion (Berg, 1983). Third, we remove the rigid circular boundary condition and treat the filament tip as a two-dimensional random walker with a linear restoring force. In this case we employ numerical random-walk simulations to determine the rate of surface-bound monomer interaction. Despite different underlying assumptions, the three approaches produce remarkably congruent results.

### Discrete random walk inside a circular boundary

Here we assume that the filament tip undergoes an unbiased random walk on a rectangular grid with a circular boundary, of radius *r*_*disc*_ (as calculated in Appendix B). We further assume that each interaction with an actin-charged polymerase delivers only one monomer to the filament. Under these conditions, the rate of surface-mediated elongation (*R*_*surf*_) does not increase linearly with soluble profilin-actin concentration but follows a saturation curve (Appendix C).

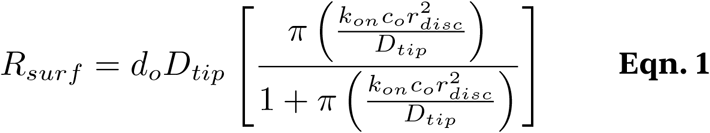

Where *D*_*tip*_ is the effective diffusion coefficient of the filament tip (Mogilner, 1996), *d*_*o*_ is the surface density of profilin-actin binding sites, and *k*_*on*_ is second-order rate constant that describes profilin-actin binding to surface-associated polymerases. This formula reproduces the observed linear dependence of filament elongation on surface density of polymerase sites (Bieling, 2018). Furthermore, at low profilin-actin concentrations (*c*_*o*_) the elongation rate becomes approximately

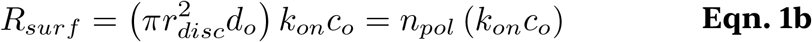

Under these conditions, a fluctuating filament tip scours polymerase-bound actin from the membrane interaction disc faster than it can be replaced from solution, and the rate-limiting step in filament elongation becomes the diffusion of profilin-actin into the membrane-interaction disc.

By contrast, high concentrations of profilin-actin rapidly refill surface-associated polymerases and the rate of filament elongation depends only on how quickly a filament tip finds the next polymerase. The elongation rate then becomes independent of both profilin-actin concentration and the size of the membrane interaction disc, asymptotically approaching a constant value. The transition between “low” and “high” concentration behaviors is driven by the

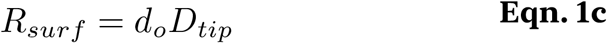

expression in parentheses in **Eqn. 1**. This unitless term, which we will call *δ*_1_, comprises the rate of polymerase loading (*k*_*on*_*c*_*o*_) multiplied by the time required for the filament tip to transit the membrane interaction disc (*r*^*2*^_*disc*_/*D*_*tip*_).

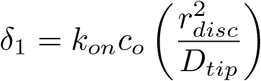

A physical interpretation of this product is that it represents the number of times an empty polymerase can be charged with profilin-actin during one sweep of the filament tip across the membrane interaction disc. Using this notation, we can rewrite **Eqn. 1** in a more compact form.

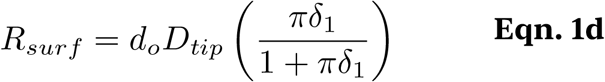

### Relative contributions of solution- and surface-mediated pathways to filament elongation

We can now quantitatively compare the rate of surface-mediated elongation to the growth of actin filaments fueled by soluble profilin-actin complexes. The Smoluchowski limit for the rate of filament elongation from solution (*R*_*soln*_) can be expressed as (Drenckhahn, 1986):

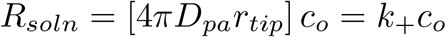

Where *D*_*pa*_ is the diffusion coefficient of profilin-actin in solution, *r*_*tip*_ is the molecular interaction radius of the filament tip, and *k*_+_ is a second-order rate constant for filament elongation. Similarly, the diffusion-limited rate of loading a surface-associated polymerase from solution can be expressed as (Berg and Purcell, 1977):

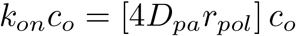

Where *r*_*pol*_ is the capture radius of the polymerase molecule. Assuming that the capture radius of a polymerase is approximately equal to that of a growing filament tip (i.e. *r*_*fil*_ ≅ *r*_*pol*_), we can use **Eqn. 1d** to describe the rate of surface-mediated elongation as a function of elongation from solution:

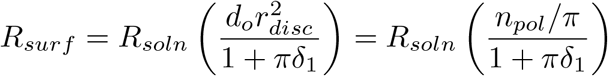

Where *n*_*pol*_ represents the average number of polymerase molecules inside the membrane interaction area. A filament growing against a polymerase-covered surface will incorporate both soluble and surface-bound monomers, so the total elongation rate will be

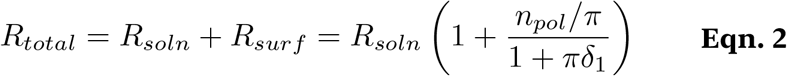

Note that the term in parentheses of **Eqn. 2** represents the fold acceleration of filament growth in proximity to surface-bound polymerases. Under these conditions the fraction of *total* actin delivered to the filament from the surface is given by the ratio:

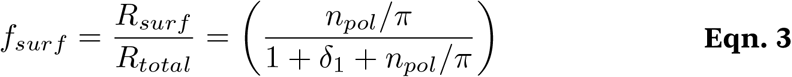

The *δ*_1_ term (see above) is linearly proportional to soluble actin concentration, so the ratio of surface-to solution-mediated elongation rates is highest at low concentrations (*δ*_1_ ≅ 0) and monotonically decreases with increasing profilin-actin concentration. We can, therefore, determine the minimum polymerase density required to match solution-mediated elongation by setting *δ*_1_ equal to zero and finding values of *n*_*pol*_ for which the expression in parentheses of **Eqn. 3** equals or exceeds one half. This occurs when

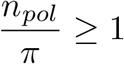

From this we conclude that, to match solution-mediated elongation, each filament must be fed by three or more surface-associated polymerases. We can also use **Eqn. 3** to calculate the effect of surface polymerases under more physiologically relevant conditions. For example, at membrane actin densities of 4000/µm^2^, surface-mediated elongation will outpace elongation from solution (i.e. *R*_*surf*_ > *R*_*soln*_ or, equivalently, *f*_*surf*_ > 0.5) at all profilin-actin concentrations below ∼250 µM. At low profilin-actin concentrations the ratio of surface-to solution-mediated assembly rates approaches a limiting value given by **Eqn. 1b**, corresponding to a *maximum* theoretical fraction of surface-derived actin of 0.88.

### Continuum approach to surface-mediated elongation

We split this calculation into two parts. Initially, we ignore the circular boundary of the membrane-interaction disc and calculate the rate at which a two-dimensionally diffusing filament tip encounters surface-bound actin (*R*_*2D*_). To calculate this rate we follow Debye and Smoluchowski (Von Smoluchowski, 1917; Debye, 1942) and treat the filament tip as a two-dimensional, absorbing disc with a size defined by the radius (*r*_*tip*_) of productive interaction with polymerase-bound actin. We then use Fick’s Laws of Diffusion to calculate the two-dimensional flux of occupied polymerase molecules into this absorbing disc (Appendix D). Our approach is valid regardless of which species —filament or polymerase— undergoes diffusive motion (Debye, 1942). Solving Fick’s equations with appropriate boundary conditions yields the following, closed-form analytical analytical expression for the rate of surface-mediated filament elongation (Appendix D):

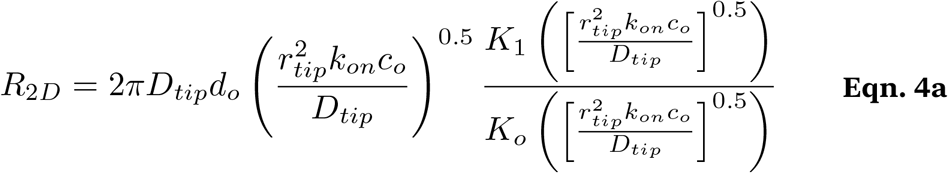

Where *K*_*o*_ and *K*_*1*_ are modified Bessel functions of order zero and one respectively. This expression is somewhat unwieldy, so we used used numerical methods to find an approximate solution that is accurate across physiologically relevant values of key parameters (i.e. *d*_*o*_, *c*_*o*_, and *D*_*tip*_). According to this analysis the interaction rate is well described by the following formula (Appendix D):

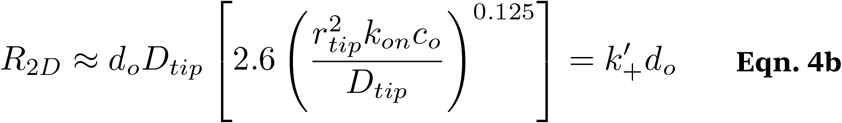

Where *k’*_*+*_ —an effective two-dimensional rate constant— has units of a diffusion coefficient (µm^2^/sec).

**Eqns 4a** and **4b** assume free diffusion of the filament tip, but bending mechanics will constrain tip motion, mostly to a circular region defined by the radius, *r*_*disc*_ (Appendix B). Within this disc, the filament visits the same polymerase molecules over and over and, when the concentration of soluble profilin-actin is low, this constraint will cause local depletion of polymerase-bound actin. Under these conditions filament elongation will be limited to the rate at which the finite number of profilin-actin binding sites (*n*_*pol*_) within the membrane-interaction disc can be refilled. Berg and Purcell (1977) calculated this rate to be

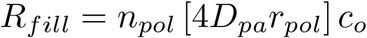

**Table C1.**
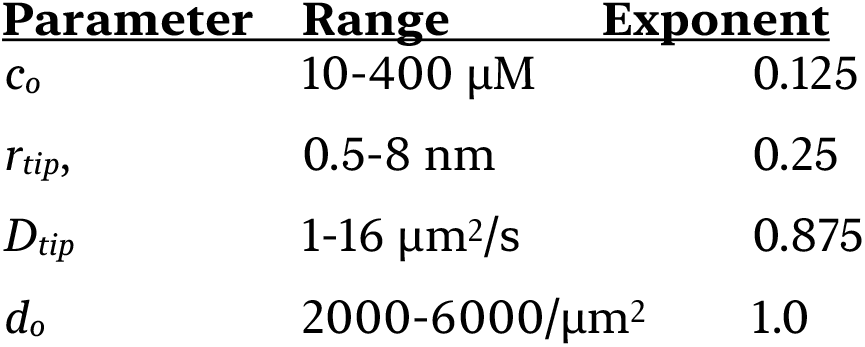
Parameter ranges and exponents.

| Table C1. Parameter ranges and exponents |  |  |
| --- | --- | --- |
| Parameter | Range | Exponent |
| $c_o$ | 10-400 $\mu\text{M}$ | 0.125 |
| $r_{tip}$ | 0.5-8 nm | 0.25 |
| $D_{tip}$ | 1-16 $\mu\text{m}^2/\text{s}$ | 0.875 |
| $d_o$ | 2000-6000/$\mu\text{m}^2$ | 1.0 |

Where *D*_*pa*_ is the diffusion coefficient of soluble profilin-actin and *r*_*pol*_ is the capture radius of a polymerase molecule. This equation holds for relatively low polymerase densities (i.e. *n*_*pol*_*r*_*pa*_<*r*_*disc*_). At higher densities (i.e. *n*_*pol*_*r*_*pa*_>*r*_*disc*_) nearby profilin-actin binding sites begin to compete with each other and the refilling rate is limited by how quickly profilin-actin diffuses into the membrane-interaction disc. Under these conditions the filling rate is independent of the polymerase density and becomes:

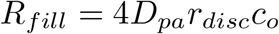

Regardless of which expression we use for *R*_*fill*_, the overall surface-associated polymerization rate (*R*_*surf*_) satisfies the expression:

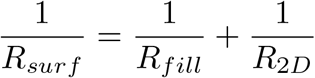

Which yields:

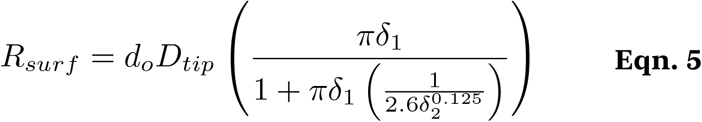

With parameters, *δ*_1_ and *δ*_2_, defined by:

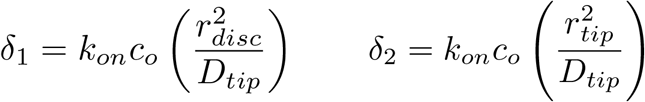

Reassuringly, **Eqn. 5** strongly resembles **Eqn. 1d**, the only difference being the presence of the *δ*_2_-containing term in the denominator of **Eqn. 5**. At low profilin-actin concentrations **Eqns. 1d** and **5** both reduce to the same form (i.e. **Eqn. 1b**). In high profilin-actin concentrations **Eqn. 5** reduces to

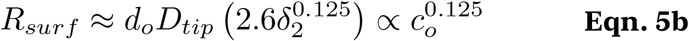

The product *d*_*o*_*D*_*tip*_ describes the filament-polymerase collision frequency but, unlike in **Eqn. 1c**, this value is not an upper bound on filament growth rate. According to **Eqn. 5b** the rate of elongation can be driven arbitrarily high (Figure 2) by increasing the profilin-actin concentration (*c*_*o*_). This difference between **Eqns. 1d** and **5** arises from a difference in underlying assumptions. The discrete random walk approach behind **Eqn. 1d** explicitly assumes that each filament-polymerase interaction transfers *at most* a single actin subunit to the growing filament. The continuum approach of **Eqn. 5**, on the other hand, implicitly assumes that a polymerase site can collect multiple actin molecules from solution and deliver them to the filament tip during one collision event. In this light we interpret the unitless expression in parentheses (*rtip*^2^*k*_*on*_*c*_*o*_/*D*_*tip*_) as the average number of profilin-actin complexes that can be loaded onto a polymerase during one filament interaction. We call this term *δ*_2_, and note its strong resemblance to the parameter *δ*_1_, described above. In physiologically relevant concentrations of profilin-actin this term (2.6*δ*2^0.125^) has a modest effect on the elongation rate (Figure 2). Specifically, the value of 2.6*δ*2^0.125^ rises from 1.0 at 50 µM profilin-actin to 1.2 at 200 µM.

**Figure 2.**
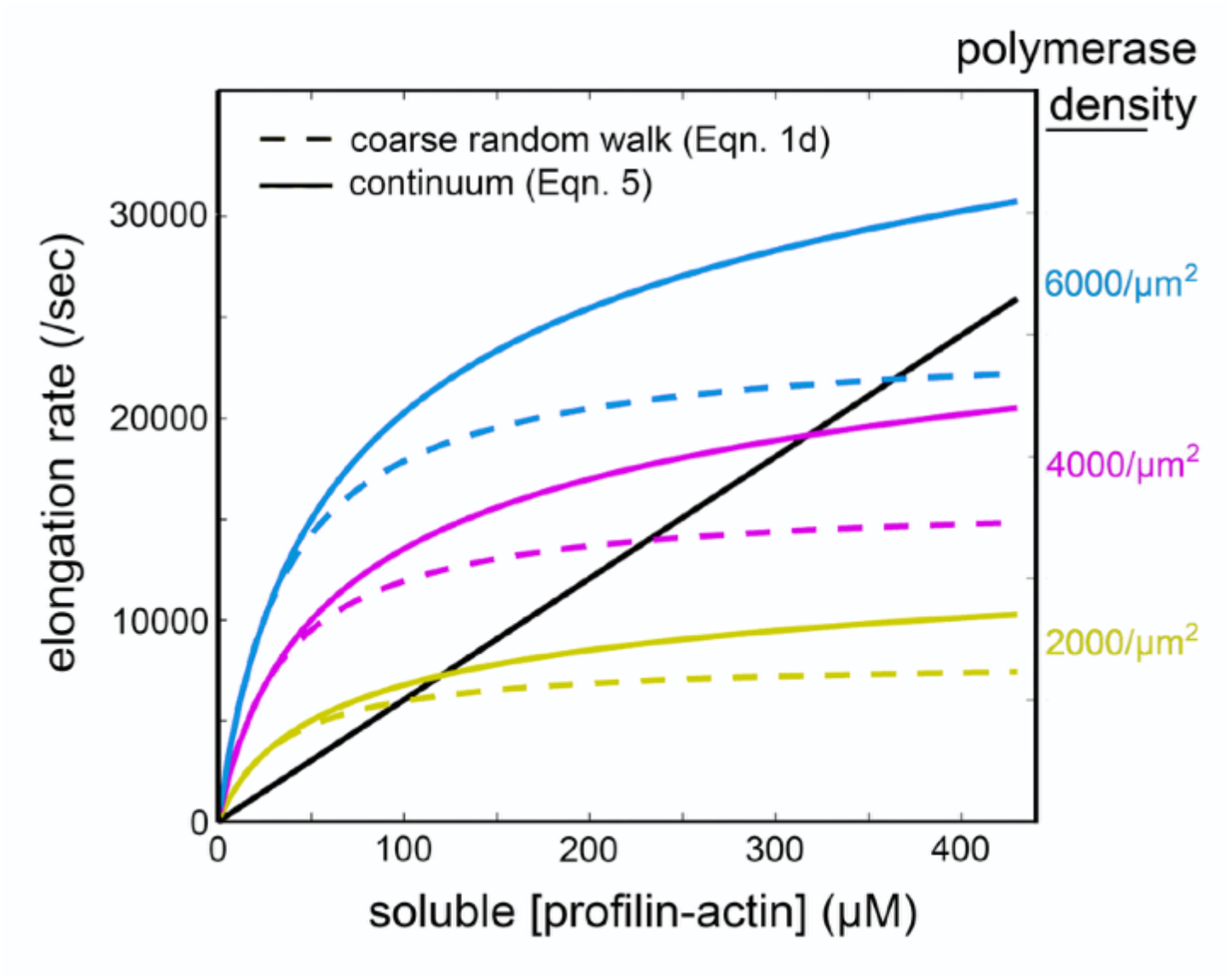
Comparison of surface-mediated elongation rates calculated using ‘coarse’ random walk (**Eqn. 1d**, dashed lines) and continuum (**Eqn. 5**, solid lines) approaches. Elongation rate as a function of soluble profilin-actin concentration was calculated at three different surface polymerase densities: 2000/µm2 (yellow), 4000/µm2 (magenta), and 6000/µm2 (cyan). The calculated rate of elongation from soluble components (black line) is shown for comparison.

### Random walk simulations of surface-mediated polymerization

To test the assumptions underlying our mathematical analysis we constructed a numerical simulation of the tip of an actin filament undergoing a constrained random walk across a polymerase-covered membrane surface (Figure 3A). Briefly, we assumed that the filament tip undergoes a two-dimensional random walk with an effective diffusion coefficient of 4 µm^2^/sec (Mogilner, 1996), while experiencing a distance-dependent restoring force directed toward the origin. We chose a spring constant that produces a root-mean-square tip displacement of 45 nm, consistent with thermal fluctuations of a 300 nm actin filament (Figure 1B, B1). We randomly distributed immobile polymerase sites at various densities across the surface, and scored an interaction every time the filament tip ventured within 2 nm (*r*_*tip*_) of an actin-charged polymerase site. To match our continuum calculations we assumed that polymerases could be reloaded while still in close proximity to filament ends, leading to the possibility of multiple elongation events per encounter. Empty polymerase sites that had given up their actin monomer to the filament were replenished at a rate proportional to the concentration of soluble profilin-actin (Berg, 1977).

**Figure 3.**
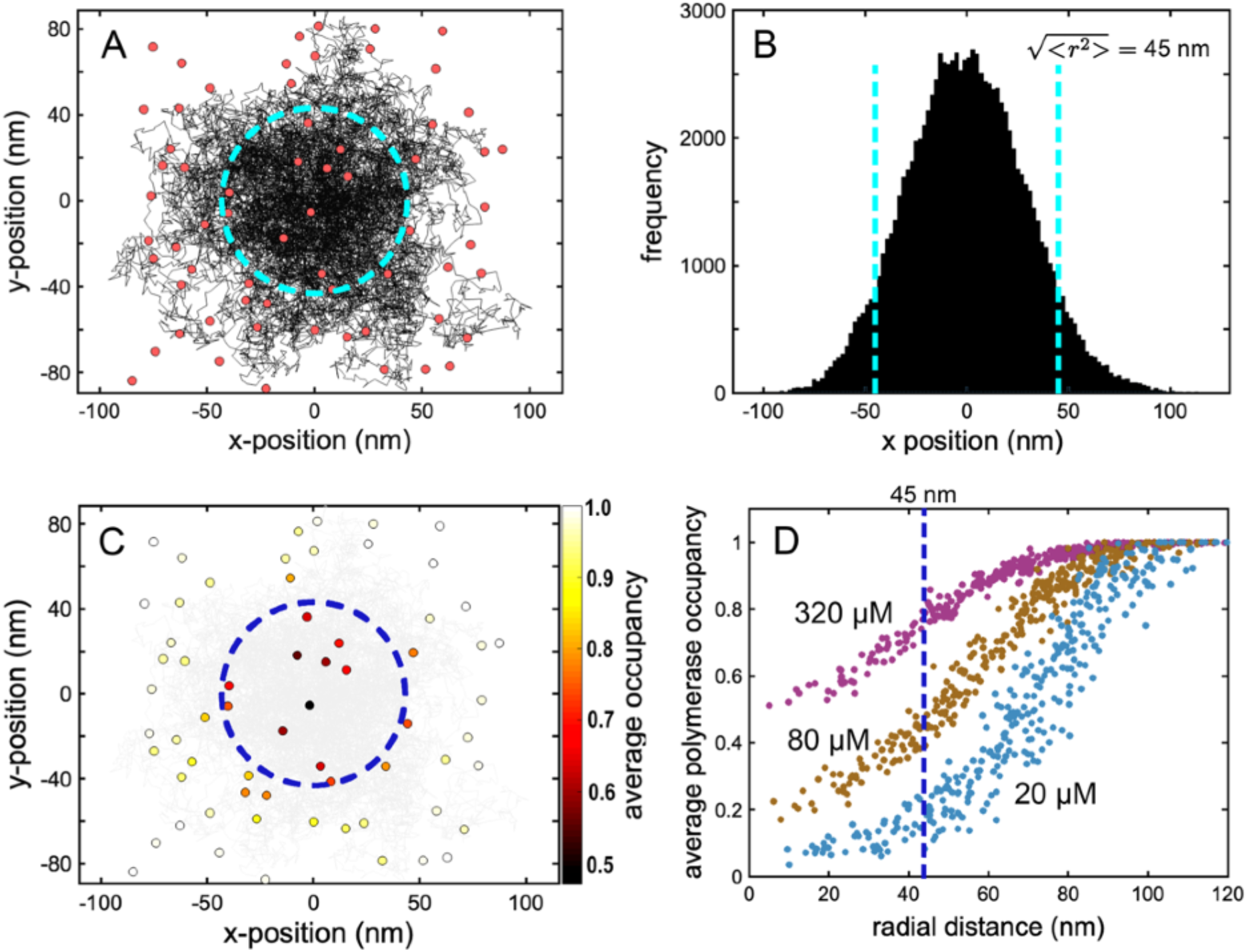
Random walk simulations of filament elongation near a polymerase-coated membrane. (A) Trajectory of a 0.02 second constrained random walk (black) across a 2D surface randomly seeded with polymerase molecules (red circles) at an equivalent of 2000/µm^2^. Dashed cyan circle: RMS radial position of the tip over the trajectory. (B) Histogram of x-projected tip positions. Dashed cyan line: RMS tip position. (C) Polymerase molecules from (A) colored according to average occupancy over a 0.1 sec trajectory. Soluble proflin-actin concentration: 320 µM. (D) Average occupancy over 0.4 second random walks at three different soluble profilin-actin concentrations: 20 µM, 80 µM, and 320 µM, as marked. Each curve is the aggregate of 5 simulations.

In our simulations the origin-directed restoring force effectively constrains trajectories to a radially symmetrical region centered on the origin (Figure 3A), and the distribution of tip positions projected onto both x-and y-axes is approximately Gaussian (Figure 3B). As predicted, the constrained nature of the random walk produces a local depletion of actin-charged polymerase molecules, primarily within the membrane interaction radius (Figure 3C), and the extent of this local actin depletion depends on the concentration of soluble profilin-actin (Figure 3D). In addition to these qualitative features, the quantitative results of random walk simulations fit remarkably well with continuum theory predictions (Figure 4). According to both theory and simulation, surface-mediated elongation “wins” at lower profilin-actin concentrations, while solution-mediated elongation wins at higher concentrations. The crossover point, where surface-and solution-mediated elongation rates are equal, increases with increasing polymerase surface density. At physiologically relevant surface densities these crossover points can significantly exceed ∼100 µM (Figure 4B) —the soluble actin concentration estimated in many cell types (see Discussion). Quantitative agreement (Figure 4B) is especially remarkable given the lack of free parameters in both the theory and simulation.

**Figure 4.**
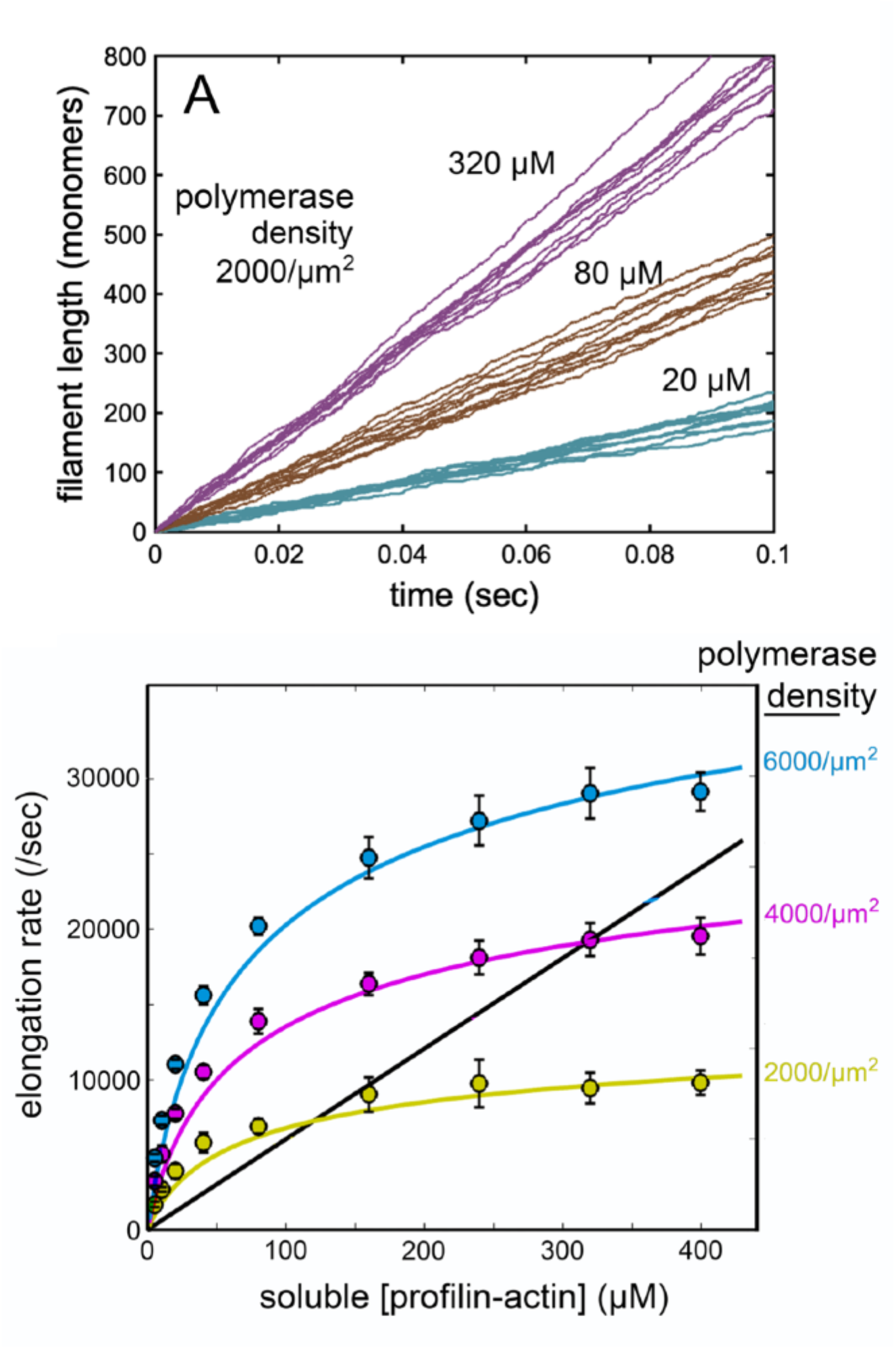
Surface-mediated elongation in random walk simulations. (A) Elongation traces from random walk simulations at three soluble profilin-actin concentrations: 20 µM, 80 µM, and 320 µM (as marked). Lines represent 10 simulations at each concentration. Polymerase surface density: 2000/µm2. (B) Comparison of elongation rates determined from simulation (color-filled circles; error bars: +/-standard deviation of 10 simulations of 0.1 seconds) and continuum theory. Color denotes polymerases density. Yellow: 2000/ µm^2^; magenta: 4000/µm^2^; blue: 6000/µm^2^. Solid curves: **Eqn. 5**. Black line: solution-mediated elongation (*R*_*soln*_).

### Effects of filament crowding

Previous experimental work revealed an inverse relationship between the local density of growing barbed ends and the elongation rate of actin filaments in a branched network (Akin, 2008; Bieling, 2018; Li, 2022). In our formalism, each filament tip is surrounded by a *radius of interference* (*r*_*inf*_) such that, when the distance between the equilibrium positions of two filament ends (*x*_*ff*_) is greater than twice this radius (i.e. *x*_*ff*_ > 2*r*_*inf*_) they will not interact with the same population of polymerases and, therefore, not affect each other’s elongation rates. Note that, because fluctuating tip positions are not uniformly distributed across the membrane interaction disc (defined by *r*_*disc*_) the value of *r*_*inf*_ will generally be larger than that of *r*_*disc*_ (Figure 5A). When two tips come close to each other (*x*_*ff*_ < 2*r*_*inf*_) the degree of interference will depend on the degree of overlap. For filaments normal to the membrane, a lens-shaped overlap region with area *A*_*OL*_, contributes a fractional overlap (*f*_*OL*_) of *A*_*OL*_/*πr*_*inf*_^2^.

**Figure 5.**
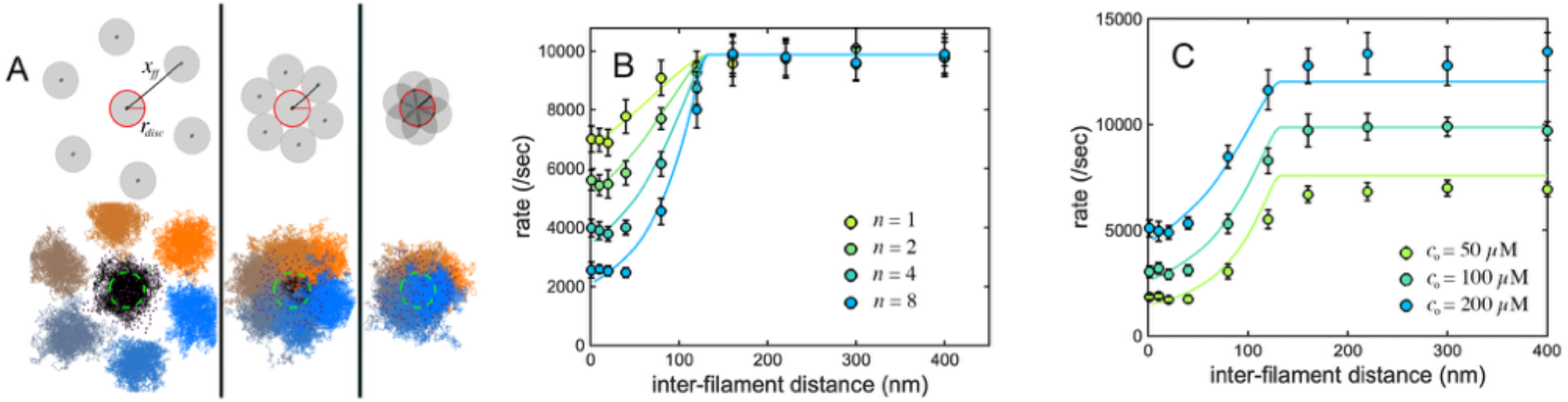
Effect of filament crowding on surface-mediated actin filament elongation. (A) Membrane interaction discs (top) and 0.1 second random walk trajectories (bottom) of seven adjacent actin filaments at three inter-filament spacings. From left to right the inter-filament spacings are: 176 nm, 88 nm, and 44 nm. (B) Effect on filament elongation rate of various numbers (*n*) of radially symmetrical, competing filaments in a single concentration of profilin-actin: *co*=100 μM. Symbols are average and standard deviation of 10 random walk simulations. Solid lines are predictions from theory (**Eqn. 6**). Different colors represent different numbers of competing filaments, as noted in the legend. (C) Effect on filament elongation rate of of *n*=6, equally spaced, radially symmetrical, competing filaments (geometry as in A) in three different profilin-actin concentrations (as noted in the figure). Symbols are average and standard deviation of 10 random walk simulations. Solid lines are predictions from theory (**Eqn. 6**). Different colors represent different profilin-actin concentrations. For B and C, the surface density of actin-binding sites (*do*) is 4000/μm^2^.

Given a filament surrounded by *n* near neighbors, each with a different fractional overlap (i.e. *f*_*OL1*_, *f*_*OL2*_, *f*_*OL3*_, …,*f*_*OLn*_), the effective number of competitors, *n*_*eff*_, is given by:

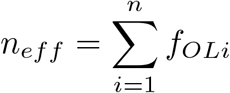

The effect of competing filaments will be to reduce the steady-state occupancy of polymerases within the membrane interaction area, and a simple modification to **Eqn. 1d** can account for this reduction:

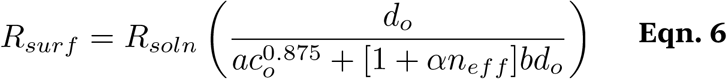

Where *α* is a fitting factor that compensates for non-uniform distribution of tip positions across *r*_*disc*_. This equation agrees well with random walk simulations (Figure 5) when the radius of interference (*r*_*inf*_) is 1.5 times *r*_*disc*_ and *α* has a value of 2. This approach can be generalized to describe different densities and distributions of membrane-proximal filament tips. This work, therefore, provides a quantitative theoretical framework for describing the effect of filament density on the rate of elongation near a polymerase-coated membrane surface and, by extension, the observed acceleration of filament growth caused by capping proteins.

### Profilin dissociation from barbed ends

Converting the actin component of a profilin-actin complex into a subunit of a growing filament requires several steps (Pollard, 1984). In the first step, a profilin-actin complex binds the filament barbed end. For a short time, the newly attached actin molecule retains its ‘monomer’ conformation (Kabsch, 1990; Schutt, 1993) and high-affinity interaction with profilin (Bieling, 2018). In less than 0.002 seconds (Funk, 2019), however, the actin shifts to a ‘filament’ conformation (Oda, 2009), and releases the bound profilin.

Because profilin blocks access to the filament barbed end, its dissociation is required for continued elongation and so the rate at which this process occurs sets an ultimate limit on the rate of filament elongation.

Note that our calculations for the effect of polymerase activity are all based on diffusion-limited encounter rates, so the fraction of monomers that enter from the solution versus the surface are not changed by the effect of profilin. That is, when a barbed end sheds a bound profilin, the rates at which it encounters a new profilin-actin complex from solution versus the surface are unchanged. **Eqn. 3**, therefore, remains valid regardless of the rate of profilin dissociation.

To account for the effect of profilin dissociation we must first convert our diffusion-limited ‘monomer-encounter’ rates (i.e. *R*_*surf*_, *R*_*soln*_, and *R*_*total*_) to elongation rates, by accounting for steric and electrostatic effects. We estimate an effective steric-and-electrostatic factor (Debye, 1942) for solution-mediated elongation, *φ*_*se*_, by dividing measured actin filament elongation rate constants (Pollard, 1986; Fujiwara, 2007) by the Smoluchowski limit calculated above (i.e. *R*_*soln*_). Assuming that steric and electrostatic effects are the same for solution- and surface-mediated polymerization (see Discussion), we apply this same correction factor to all of our calculated rates. The profilin dissociation-limited elongation rates (*E*_*soln-p*_, *E*_*surf-p*_, and *E*_*total-p*_) are now given by:

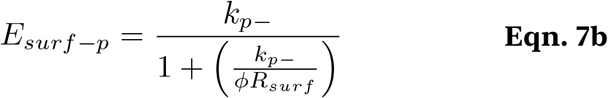

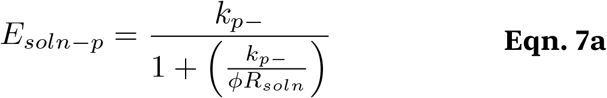

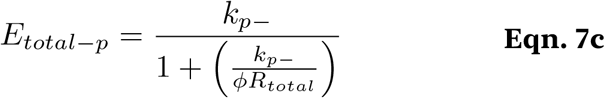

Where *k*_*p-*_ —the rate constant governing profilin dissociation— has a measured value of ∼550/s, equivalent to the rate of solution-mediated filament elongation in the presence of 55 µM soluble actin. This means that, at cellular profilin-actin concentrations of ∼100 µM, filament elongation will occur at a more or less fixed rate, regardless of the influence of surface-associated polymerases (Figure 6A). Note that, even though the polymerase-coated surface may not increase the elongation rate, more subunits may still be delivered to the filament from the surface than from solution (**Eqn. 3**).

**Figure 6.**
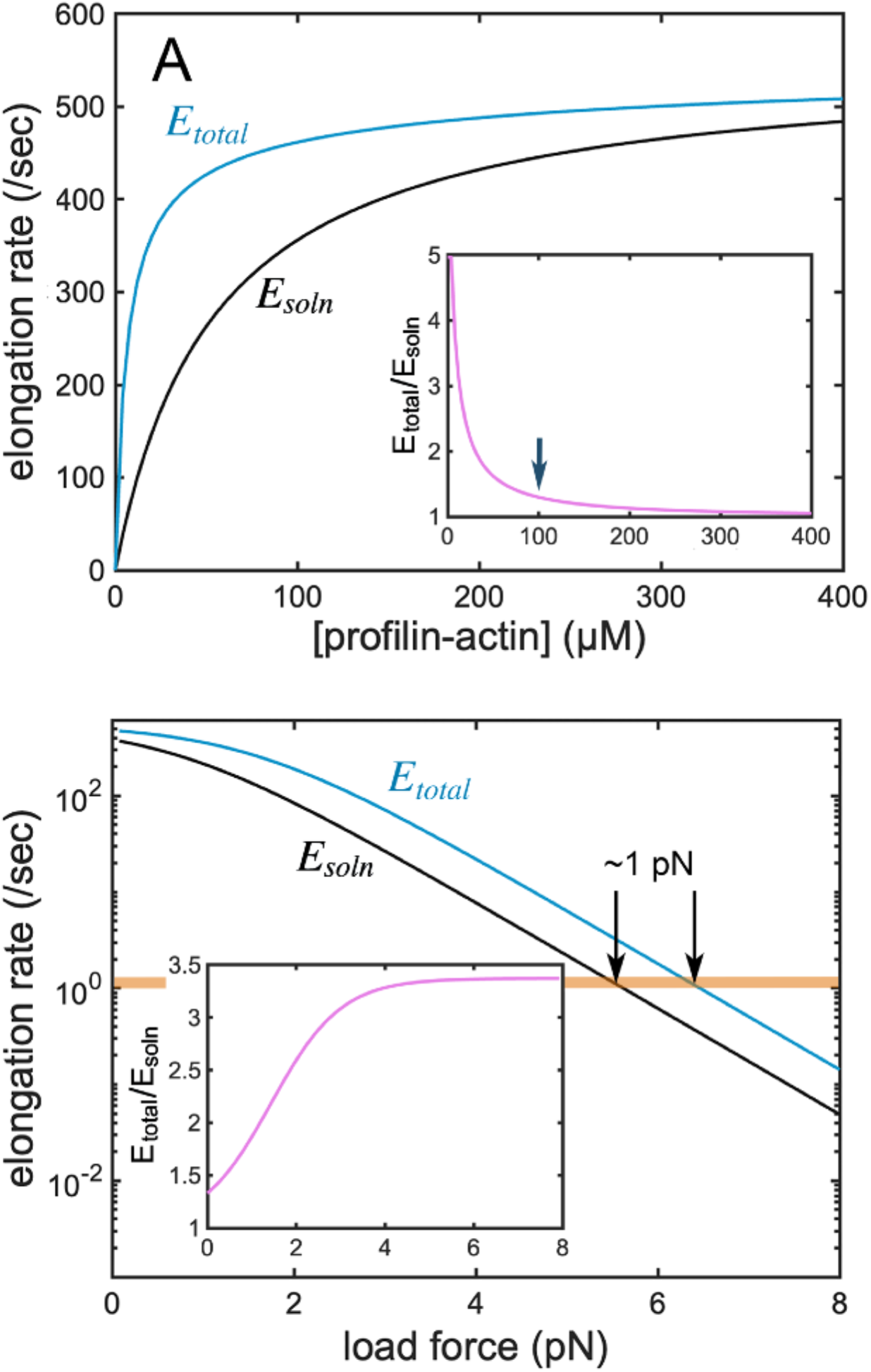
The effects of profilin dissociation and applied force on surface-mediated filament elongation. (A) Filament elongation limited by the rate of profilin dissociation. Black: solution-mediated elongation; blue: solution-plus surface-mediated elongation (*d*_*o*_=4000/µm^2^). Inset: rate enhancement due to polymerase activity. X-axis same as main plot. Arrow marks co=100 µM. (B) Filament growth against a load force. Black: solution-mediated elongation; blue: total (solution-plus surface) elongation (*c*_*o*_=100 µM; *d*_*o*_=4000/ µm^2^). Arrows mark intersection of elongation rate with monomer dissociation rate. Inset: rate enhancement due to polymerase activity. X-axis same as main plot.

If surface-associated polymerases cannot accelerate filament elongation beyond a speed limit imposed by profilin dissociation, do they have any effect on the rate of network growth? In soluble actin concentrations of ∼100 µM and zero load force, the answer appears to be: “not much.” When forces are applied to the network, however, the answer is very different.

### The effect of force

When an actin filament grows against an opposing force the elongation rate slows by an amount that reflects the energy required to insert each new subunit against the applied force. Specifically, if subunit addition extends the filament length by *δ*, against opposing force *f*_*o*_, elongation slows exponentially, by a factor equal to:

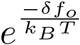

Where *k*_*B*_ is Boltzmann’s constant and *T* is absolute temperature. This factor follows directly from thermodynamic analysis (Hill, 1982), while Elastic Brownian Ratchet (Mogilner, 1996) theory provides a clear microscopic rationale. Briefly, this factor represents the fraction of exponentially distributed filament fluctuations that are large enough to permit insertion of a *δ-*sized building block between the filament tip and the load surface (Mogilner, 1996). At room temperature, *k*_*B*_*T* is approximately 4.11 pN-nm, and adding a profilin-actin complex to the barbed end of a filament increases its length by ∼5.2 nm (i.e. the 2.7 nm actin subunit overlap plus the 2.5 nm length of a bound profilin). The ratio, *δ/ k*_*B*_*T*, therefore, becomes ∼1.27/pN. If we assume that force has little or no effect on the rate of conformational change in the terminal monomer or the dissociation of profilin (Mogilner, 1996), the rates of elongation become:

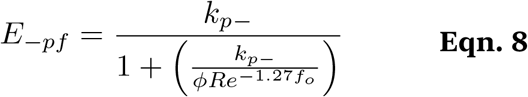

Assuming realistic values for c_o_ (100 µM) and d_o_ (4000/µm^2^) we can plot the adjusted rates of surface- and solution-mediated elongation as well as the total elongation rate (*E*_*surf-pf*_, *E*_*soln-pf*_, and *E*_*total-pf*_), as functions of applied force (Figure 6B). Under these conditions the surface-polymerases have a modest (1.3-fold) effect on zero-load elongation. As force increases and elongation rates decrease, however, the effect of the polymerases becomes more pronounced. For forces in the region of 4-7 pN (the estimated range of stall forces for single actin filaments at the leading edge of crawling cells) the polymerase-coated surface enhances elongation by ∼3.5-fold (Figure 6B). This increased enhancement reflects a shift from a regime in which elongation is limited by the rate of profilin dissociation to a regime in which elongation is limited the rate of monomer encounter. In other words, membrane-associated polymerases provide no advantage when profilin dissociation limits elongation, but accelerate growth under conditions where monomer encounter is rate-limiting.

Note that when elongation slows to the point where it equals the rate of monomer dissociation (∼1/sec) filament growth will stall. For the conditions shown in Figure 6B, surface polymerases increase this force by ∼1 pN (∼18%).

## Discussion

Similar to Edwin Abbott’s, “Flatland” (Abbot, 1884), actin filament assembly is a “romance of many dimensions.” Soluble profilin-actin complexes, diffusing in three-dimensions, are collected onto a two-dimensional membrane surface by polymerase molecules that facilitate their conversion into one-dimensional filaments. The proline-rich profilin-actin binding sites found on WAVE/WASP family proteins function as the major inter-dimensional portal for assembly of branched actin networks.

Experimentally, the actin polymerase activity of surface-bound WAVE/WASP proteins exhibits two key features: (i) it is highly distributive and (ii) it increases linearly with surface density (Bieling, 2018). The distributive nature of the activity means that a given WAVE/WASP molecule spends very little time bound to the end of a growing filament, while the density dependence indicates that the elongation rate depends on how quickly a filament tip encounters fresh, actin-charged polymerases. In these chance encounters, who finds whom? Do mobile polymerase molecules find a more or less static filament, or does a dynamic filament tip search for nearby polymerases? Our analysis assumes that tip mobility drives the interaction, but our mathematical formalism can also account for polymerase mobility. Similar to the Debye-Smoluchowski equation, we could replace the *D*_*tip*_ term in **Eqn. 1** with (*D*_*tip*_ + *D*_*pol*_), where *D*_*pol*_ represents the diffusion coefficient of the polymerase molecules. We note that polymerase molecules may not undergo long-range motion, but flexible, C-terminal sequences may still permit significant segmental motion of bound profilin-actin complexes. We could account for this by adding an effective interaction radius, *r*_*pa*_, for the polymerase-bound actin to **Eqn. 1**, replacing *r*_*tip*_ by (*r*_*tip*_ + *r*_*pa*_).

Two-dimensional diffusion of a protein in a lipid membrane is generally much slower than three-dimensional diffusion of a soluble protein (McCloskey, 1986), which is one reason we ignore the lateral diffusion of WAVE molecules in our model. Interestingly, the effective diffusion coefficient calculated by Mogilner and Oster (1996) for a thermally fluctuating filament tip (4 µm^2^/sec) is remarkably similar to the measured diffusion coefficient of profilin-actin in cytoplasm (5 µm^2^/ sec). Also, at the estimated membrane densities of WAVE/WASP family proteins, the intermolecular spacing of surface-associated actin binding sites (<22 nm) is closer than of actin molecules in a 100 µM solution. These facts, together with the higher probabilities of molecular interaction in two dimensions (Huang, 2024) help explain the ability of clustered WAVE/WASP molecules to accelerate filament assembly, even when the soluble actin concentration is >100 µM.

We assume that steric and electrostatic effects are the same for solution- and surface-mediated elongation. We take this approach partly to demonstrate that invoking these effects is not required to explain the enhancement of polymerization caused by clustering WAVE/WASP proteins on a membrane surface. Formally, the steric factor could either: (i) decrease due to a decreased rotational diffusion coefficient of the tethered profilin-actin (Northrup, 1992); (ii) increase due to reduced dimensionality and increased search efficiency of the tethered species (Windisch, 2006); or (iii) remain unchanged. When future biophysical experiments and/or molecular dynamics simulations provide additional data relevant to this question, we can modify the value of the steric/ electrostatic factor in **Eqns 7a, 7b, 7c**, and **8**.

In mammalian cells the estimated concentrations of monomeric actin near sites of branched network assembly vary considerably. Many estimates fall in the range of ∼100 µM (Mogilner, 2002; Kiuchi, 2011) to 150µM (Koestler, 2009), while some fall as low as 10µM (Novak, 2008) and some as high as 250 µM (Fox, 1993). Our theory and simulations demonstrate that surface-associated polymerases effectively accelerate filament elongation across this entire range of soluble actin concentrations. At low concentrations the rate of surface-mediated polymerization is limited by how quickly the small number of polymerases that interact with each filament (*n*_*pol*_) can be replenished from solution. For a diffusion-limited interaction, this rate depends linearly on the concentration (*c*_*o*_) and diffusion coefficient (*D*_*pa*_) of the soluble profilin-actin complexes. At high concentrations of soluble actin, however, the rate is limited by two linked processes, each with its own (effective) diffusion coefficient. The elongation rate is still influenced by the re-charging of depleted polymerases, but it also depends on the rate at which fluctuating filament tips encounter actin-charged polymerases.

Our formalism recaptures key features of previously published experimental data. Specifically, Bieling (2018) found that the rate of surface-mediated polymerization increases linearly from 0 to 2.75-times the rate of solution-mediated elongation as WAVE1 surface density increases from 0 to 1850/µm^2^. These experiments were performed in the presence of 1 µM soluble profilin-actin. Given this low actin concentration and relatively low WAVE1 surface density, the ratio of surface- to solution-mediated elongation rates will be described by the limiting case of **Eqn. 1b**. Assuming that the filament tip and polymerase molecule have similar interaction radii (i.e. *r*_*tip*_∼*r*_*pa*_), then *R*_*surf*_/*R*_*soln*_= 2.75 = *n*_*pol*_/*π*. From this, we estimate that each filament tip interacts with an average of 8.6 actin binding sites. If each WAVE1 molecule binds two profilin-actin complexes, then on average each filament is fed by 4.3 WAVE1 molecules. From this, we estimate that the surface interaction area of these filaments is ∼2300 nm^2^. This is approximately 36% of the interaction area of our model 300 nm filament oriented normal to a membrane surface (Appendix B). The smaller interaction area likely reflects the significant difference in geometry. Specifically, the filaments in the Bieling experiments elongated parallel to WAVE1-coated surfaces, so the interaction area traced by their fluctuating tips was most likely a “croissant” shape, rather than a radially symmetrical disc. In addition, these filaments were typically several microns long and the distance between their growing barbed ends and the nearest point of surface tethering was likely longer than 300 nm, leading to larger RMS tip deviations.

Finally, quantitative results of this theory are consistent with our recent measurements of the fraction of actin delivered to leading-edge lamellipodial networks by surface-associated polymerases that contain profilin-actin binding sites (Skruber, 2024). This experimental study reported an approximate 75% reduction of the incorporation rate of covalently linked actin-profilin complexes when a point mutation (H133S) was introduced into the profilin moiety that reduced affinity for proline-rich binding sites found in surface-associated polymerase molecules. This result implies that ∼75% of the actin enters the network from the membrane surface and it agrees well with our theory, which predicts 75% surface contribution at soluble profilin-actin concentrations of ∼100 µM and surface actin densities of ∼4000/µm^2^.

## Acknowledgments

This work was supported by grants from the National Institutes of Health (1R35 GM118119) and the Howard Hughes Medical Institute. We are grateful to members of the Mullins Lab for helpful discussions and critical readings of the manuscript, especially Sam Lord and Arthur Charles-Orszag. We are also indebted to Alex Mogilner, who suggested that we perform the discrete random walk simulation, and to Rob Phillips and Tom Pollard for critical readings of early drafts of this manuscript which improved both the argument and presentation.

## Appendix A

### List of variables and parameters

*λ*_*p-*_: Persistence Length of an actin filament Length of actin filament
*D*_*pa*_: Diffusion coefficient of soluble profilin-actin complexes
*D*_*tip*_: Effective diffusion coefficient of thermally fluctuating actin filament tip
*r*_*disc*_: Root Mean Square (RMS) deflection of membrane-adjacent filament tip
*r*_*tip*_: Capture radius of the barbed end of an actin filament
*r*_*pol*_: Effective profilin-actin capture radius of a surface-bound polymerase
*n*_*pol*_: Number of polymerase molecules inside a membrane-interaction region
*c*_*o*_: Actin monomer concentration
*k*_*on*_*c*_*o*_: Rate of loading profilin-actin onto a surface-associated polymerase site
*d*_*o*_: Total surface density of profilin-actin binding sites (/µm^2^)
*d*_*pa*_: Surface density of occupied profilin-actin binding sites (/µm^2^)
*R*_*soln*_: Rate of soluble profilin-actin monomers encountering a filament tip
*R*_2*D*_: Rate of surface-associated profilin-actin encountering a nearby filament tip
*R*_*disc*_: Rate of soluble monomers encountering the membrane-interaction disc
*R*_*surf*_: Rate of surface-associated profilin-actin encountering a nearby filament tip
*E*_*soln*_: Filament elongation rate from soluble profilin-actin monomers
*E*_*surf*_: Filament elongation rate from surface-associated profilin-actin
*E*_*total*_: Filament elongation rate from soluble *and* surface-associated profilin-actin
*F*_*surf*_: Fraction of filamentous actin incorporated from surface polymerases
*f*_*OL*_: Fractional overlap of membrane-interaction regions of adjacent filaments
*n*_*eff*_: Effective number of competing filaments in a membrane interaction region
*x*_*ff*_: Distance between the equilibrium positions of two filaments tips

## Appendix B

### Size of the *membrane interaction disc* traced by fluctuating filament tips

Filaments in a functional branched actin network generally range between 150-800 nm in length (Svitkina, 1999; Akin, 2008; Koestler, 2009), much shorter than their mechanical persistence length (*λ*_*p*_). At these lengths, a filament behaves as a relatively stiff beam (Fig. B1). If we consider only small-amplitude, planar bends, deflection of the filament tip follows Hooke’s Law, meaning that displacement from equilibrium (*r*) is directly proportional to applied force (*F*):

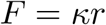

Where the effective spring constant (*κ*) depends on the persistence length (*λ*_*p*_) and actual filament length (*L*):

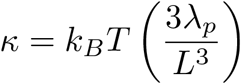

Where *T* is absolute temperature and *k*_*B*_ Boltzman’s constant. From this, we can calculate the energy stored in the bent polymer:

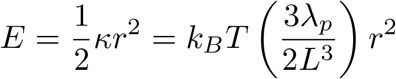

If the average energy stored the bent polymer is *k*_*B*_*T*, this yields:

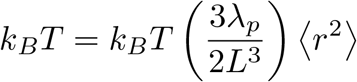

And the root-mean square (RMS) deviation of the tip is:

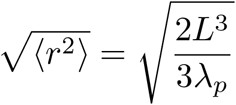

Assuming an average filament length (*L*_*av*_) of ∼110 promoters (∼300 nm), measured from the free barbed end to the network-anchored pointed end, the root-mean square (RMS) displacement of the tip will be ∼45 nm.

**Figure B1.**
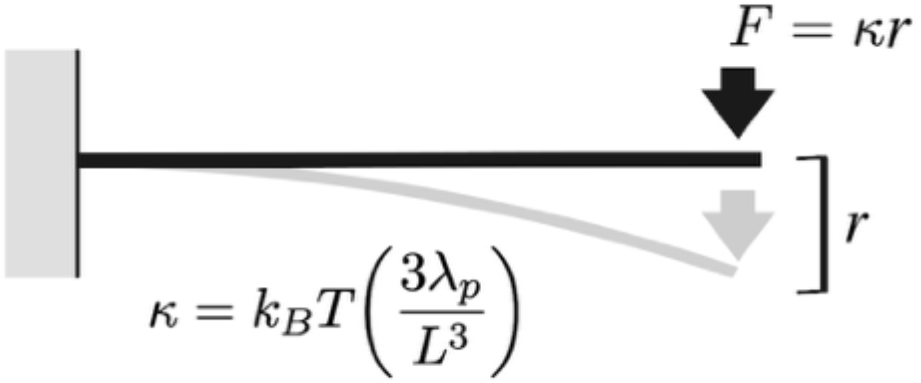
Small deflection bending of a stiff beam. The deflection is proportional to applied force, with an effective spring constant that depends on beam length and persistence length.

## Appendix C

### Analytical solution of a filament tip interacting with surface-bound polymerases via a simple, constrained random walk

Here we calculate the rate at which a filament tip undergoing a simple random walk, confined to a circular region of radius *r*_*disc*_ (calculated in Appendix B), interacts with randomly dispersed polymerase molecules. By “simple” random walk we mean that step direction does not depend on tip position (i.e. is not influenced by an elastic restoring force). We approximate the polymerase molecules as squares of length and width, *a*. (Figure C1). The number of actin-bound (occupied) polymerases inside the interaction disc is *n*_*op*_, and the number of unoccupied polymerases is *n*_*up*_. The fraction of occupied polymerases within the disc is, therefore,

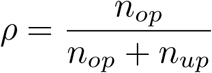

Surface-bound polymerases bind soluble profilin-actin complexes (concentration - *c*_*o*_) at a rate of *k*_*on*_*c*_*o*_, where *k*_*on*_ is an effective rate constant whose value depends on the sizes of the molecules and the diffusion coefficient of the soluble profilin-actin. Under these conditions the total rate of profilin-actin uptake by unoccupied polymerases in the interaction disc is,

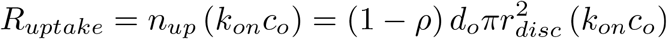

where *d*_*o*_ is the surface density of WAVE/WASP-family polymerases. Note that, at steady state, the rate of surface-mediated filament elongation (*R*_*surf*_) matches the total rate of profilin-actin uptake into the membrane interaction disc (*R*_*uptake*_). The motion of the filament tip is characterized by an effective diffusion coefficient, *D*_*tip*_ (Mogilner, 1996). We can approximate this by a discrete, two-dimensional random walk with step size *a* (the width of a polymerase molecule) and stepping rate *k*, given by

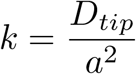

**Figure C1.**
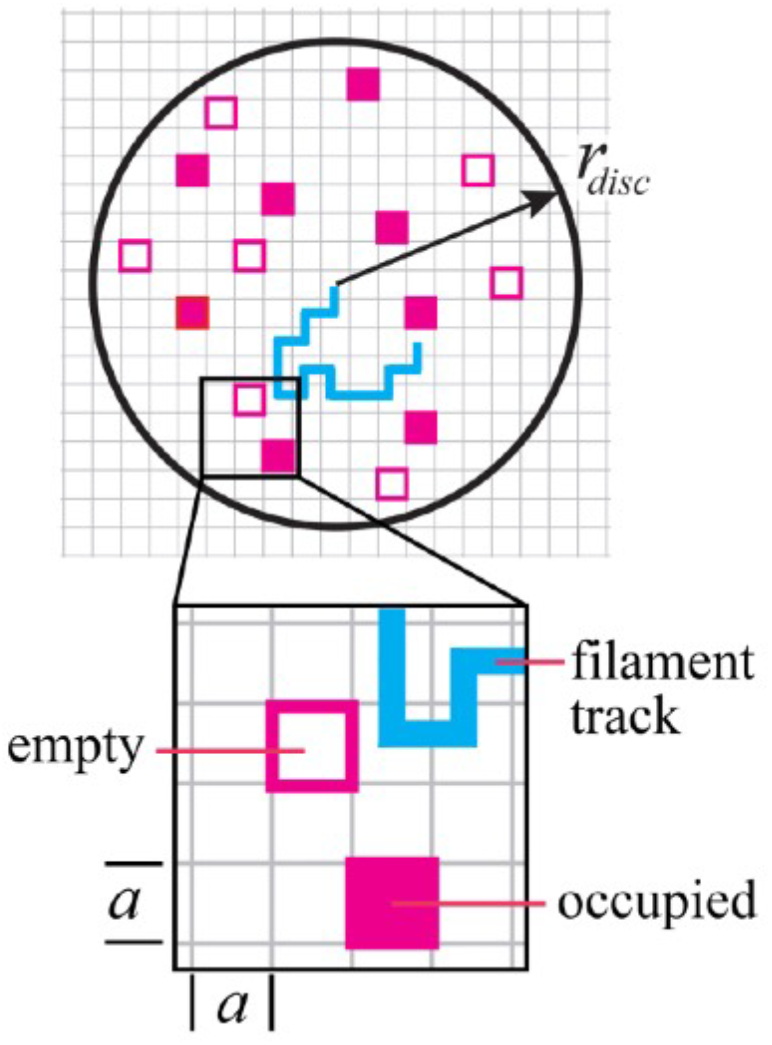
Top: filament tip (blue) doing a random walk on a circular membrane-interaction disc (radius -*r*_*disc*_) filled with polymerase molecules (red squares). Open squares are unoccupied; filled squares are charged with profilin-actin. Bottom: zoom.

The rate of filament elongation is now given by the stepping rate (k) multiplied by the probability that the destination square contains a profilin-actin-charged polymerase (*P*_*occupied*_). To calculate the occupancy probability we divide the number of charged polymerases in the interaction disc by the total number of polymerases that can be packed into the disc

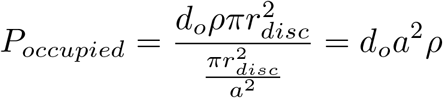

And the rate of elongation becomes (**Eq. C1**)

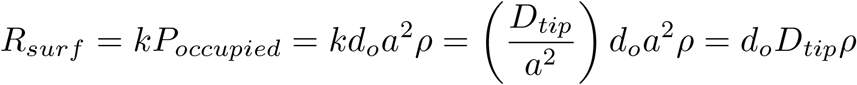

At steady state, the rate of elongation matches the rate of profilin-actin uptake into the interaction disc, such that

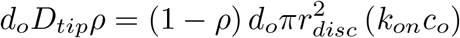

Solving for the fraction of occupied polymerases (*ρ*) yields

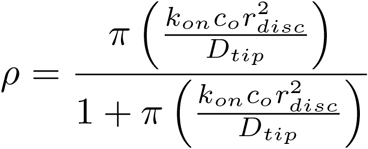

Finally, substituting this expression for *ρ* into **Eq. C1** produces an equation for the elongation rate as a function of the soluble profilin-actin concentration (**Eq. C2**)

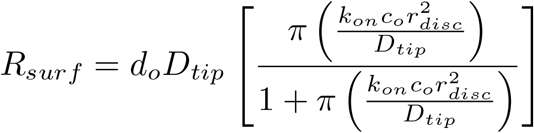

Note that the term in parentheses is unitless. The value of r2disc/Dtip is proportional to the time required for the fluctuating tip to traverse the interaction disc while konco is the rate of profilin-actin delivery to a polymerase molecule. The product of these two terms, therefore, can be interpreted as the number profilin-actin complexes that can be delivered to each polymerase during one sweep of the filament across the disc.

## Appendix D

### Numerical solution of Fick’s equation in two dimensions with polymerase refilling from solution

In this section we calculate a two-dimensional ‘Smoluchowski limit’ on the rate at which a growing filament tip interacts with profilin-actin-charged polymerase molecules on a membrane surface. We treat the filament tip as a circular hole whose radius (*r*_*tip*_) represents the effective radius of interaction with polymerase-bound actin. For boundary conditions, we assume that the density of actin-bound polymerases is zero at the edge of the filament (*d*_*pa*_(*r*_*tip*_) = 0), and asymptotically approaches the total polymerase surface density (*d*_*o*_) at long distances (*lim*_*r*→∞_[*d*_*pa*_(*r*)] = *d*_*o*_). With these boundary conditions, we use Fick’s Second Law of Diffusion to calculate the gradient of occupied polymerase molecules around the filament tip (Von Smoluchowski, 1917; Debye, 1942). Note that this approach is valid regardless of which species —filament or polymerase— undergoes diffusive motion (Debye, 1942).

If we neglect refilling of depleted polymerases by soluble profilin-actin complexes, and assume a radially symmetrical distribution of actin-bound polymerases around the filament tip, the steady-state form of Fick’s Second Law in two dimensions is:

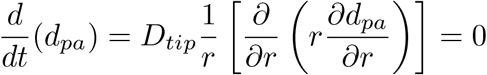

Where *D*_*tip*_ is the diffusion coefficient of the filament tip skating across the membrane surface. Unlike the three-dimensional case, this partial differential equation has no solution that satisfies the given boundary conditions. If, however, we assume that unoccupied polymerase sites are refilled from a constant reservoir of soluble profilin-actin, the steady-state distribution of occupied polymerases must satisfy:

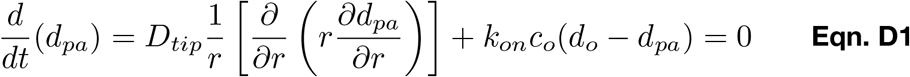

The additional term in this equation represents the rate at which unoccupied polymerases are recharged from solution. This depends on the density of unoccupied polymerases (*d*_*o*_*-d*_*pa*_), the solution concentration of profilin-actin complexes (*c*_*o*_), and a rate constant (*k*_*on*_) for binding of soluble proteins to membrane-associated sites (Berg, 1977).

To solve this equation for the surface density of loaded polymerases (*d*_*pa*_) we first make the following three substitutions:

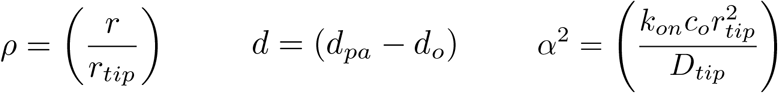

With these substitutions, we can rearrange **Eqn. D1** into the following form:

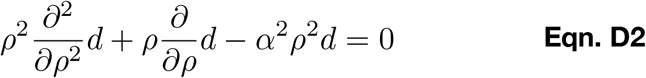

This rearranged equation has the form of a modified Bessel’s Equation of order zero, whose solution is given by a linear combination of the zero-order modified Bessel functions, *I*_*o*_ and *K*_*o*_.

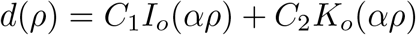

Where *C*_*1*_ and *C*_*2*_ are constants. We determine the values of *C*_*1*_ and *C*_*2*_ by applying the boundary conditions. The function *I*_*o*_ grows without bound, so the condition that *d* is bounded requires *C*_*1*_=0. When *r* = *r*_*tip*_, the substituted parameter *γ* = 1. The boundary condition that *d*_*pa*_(*r*_*tip*_) = 0, therefore, implies that

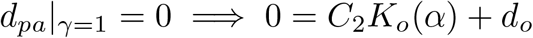

And

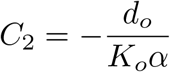

Using this value for *C*_*2*_ and substituting for *d* and *ρ*, we obtain a closed-form solution for the steady-state distribution of actin-charged polymerases

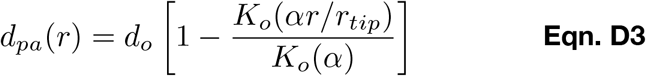

We can use this formula, together with Fick’s first law of diffusion, to compute the rate (*R*_*2D*_) at which the filament tip interacts with surface-associated actin monomers. We simply multiply the diffusion coefficient of the filament tip (*D*_*tip*_), the density gradient at the filament edge 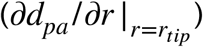, and the circumference of the monomer binding site on the end of the filament (2*πr*_*tip*_).

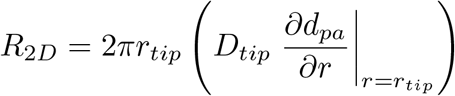

Substituting **Eqn. D3** into the above expression yields

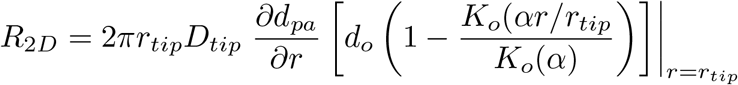

Which becomes

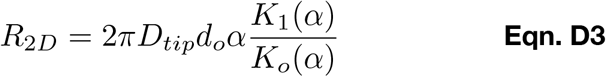

Where *K*_*1*_ is a modified Bessel function of order one. Note that the dimensionless parameter *α*^2^ resembles the previously defined parameter, *δ*_1_

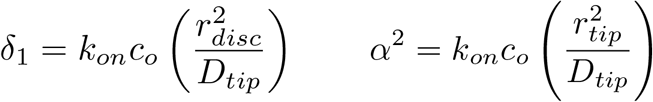

Because of this similarity, we define a new parameter, *δ*_2_ = *α*^2^. A physical interpretation of *δ*_1_ is that it represents the number of times one polymerase molecule can be loaded from solution with profilin-actin during the time it takes the filament tip to traverse the entire membrane interaction disc. Similarly, *δ*_2_ can be understood as the number of profilin-actin complexes that can bind a polymerase molecule during the course of one filament-polymerase interaction. It makes intuitive sense that the rate membrane-dependent filament elongation would depend on these parameters.

To validate our analytical solution and find a simpler, approximate expression for *R*_*2D*_, we performed a numerical simulation of two-dimensional diffusion of ligands into an absorbing disc located at the origin. Briefly, we modeled diffusion in radial coordinates on a set of concentric rings, each with a fixed width (*δ*). Given values for *δ* and filament tip diffusion coefficient (*D*_*tip*_) we chose a time step (*τ*) such that *τ* = *δ*^2^/*D*_*tip*_. We set up an initial, uniform distribution of loaded polymerases, with *d*_*pa*_ = *d*_*o*_ for all *r* > *r*_*tip*_, and stable boundary conditions of *d*_*pa*_ = 0 at *r* = *r*_*tip*_ and *d*_*pa*_ = *d*_*o*_ at *r* > 1000*δ*. At each time step, *τ*, we assumed that the molecules in each concentric ring move inward or outward with the ratio of in-to-out determined by the areas of the adjacent rings. We then calculated the number of profilin-actin complexes recruited to each ring from solution by multiplying the area of the ring by the density of unoccupied polymerases, the solution concentration of profilin-actin, the rate constant for polymerase charging and the time increment: *k*_*on*_*c*_*o*_(*d*_*o*_-*d*_*pa*_)*τ*. We obtained the steady-state distribution of actin-charged polymerases by iterating time steps until the change in distribution became negligible. We validated this approach by simulating diffusion into a three-dimensional, spherical absorber and comparing the results to von Smoluchowski (1917).

Our simulations produced stable distributions of actin-charged polymerases around the filament tip that matched the distributions calculated from our analytical solution, **Eqn. D4** (Figure D1). We used these distributions to calculate the flux of actin into the filament (*R*_*2D*_) at different values of the relevant parameters (*k*_*on*_*c*_*o*_, *r*_*tip*_, *D*_*tip*_, and *d*_*o*_).

**Figure D1.**
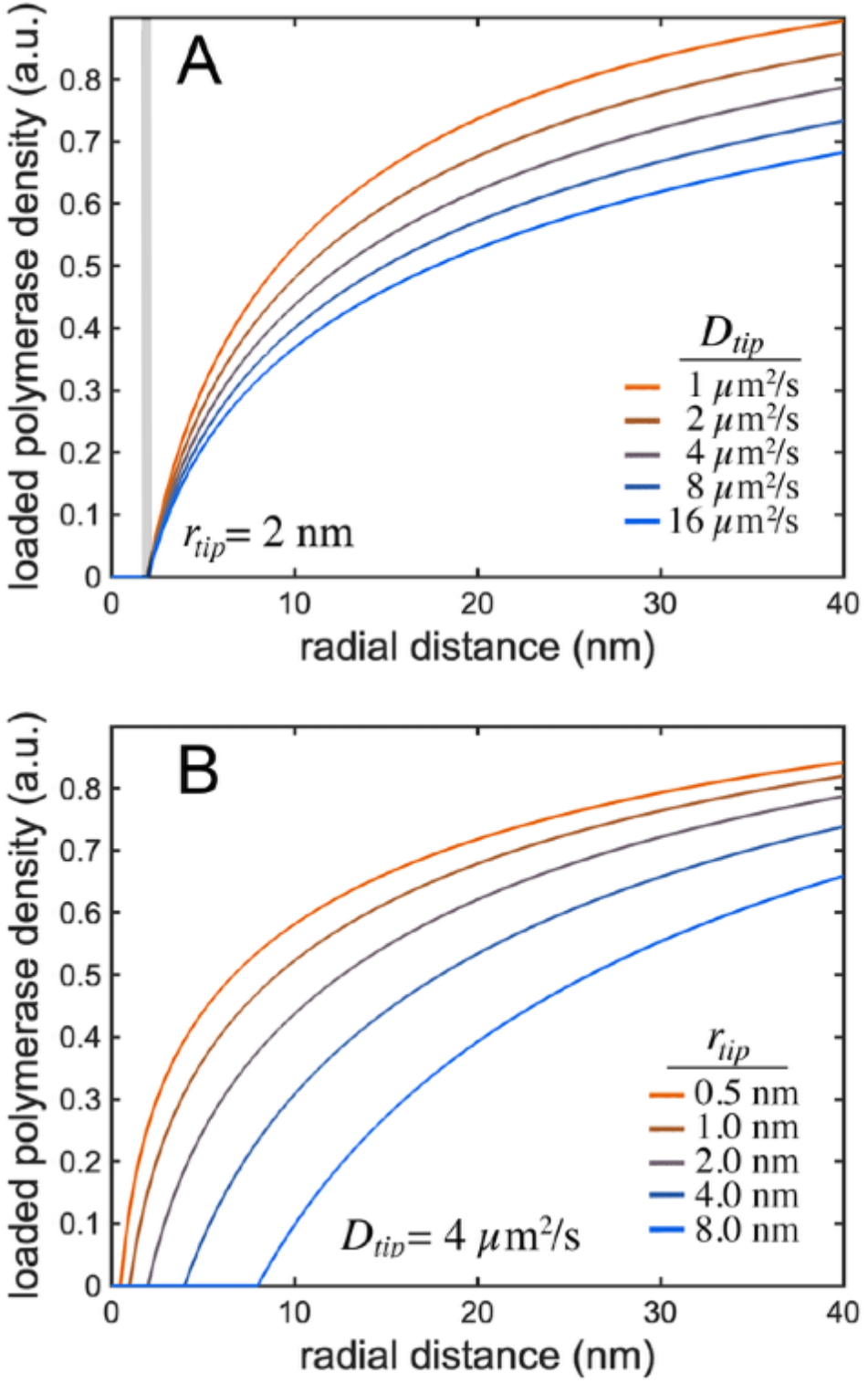
Steady-state distributions of actin-charged polymerases around an absorbing filament tip, determined by numerical simulation with various parameter values: (A) varying diffusion coefficients and (B) varying capture radii of the filament tip. For all conditions, *c*_*o*_=100µM.

Across physiologically relevant values of each parameter (Table D1), the interaction rate was well approximated as a simple power law (Figure D2, Table D1). Combining the dependences on all of the parameters into a single equation yields:

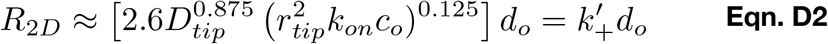

The above equation can be rearranged into a slightly more interpretable form:

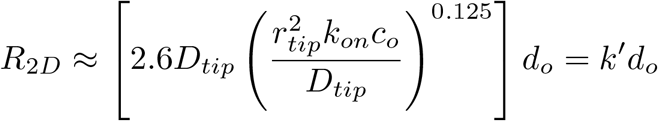

The product of diffusion coefficient and surface actin density (*D*_*tip*_*d*_*o*_) is proportional to the rate of collision between a polymerase and the filament tip, while the unitless term in parentheses (defined above as *δ*_2_) can be interpreted as the number of times a polymerase can be loaded with profilin-actin from solution during a single collision event. The term in square brackets has the form of a rate constant (*k’*).

**Figure D2.**
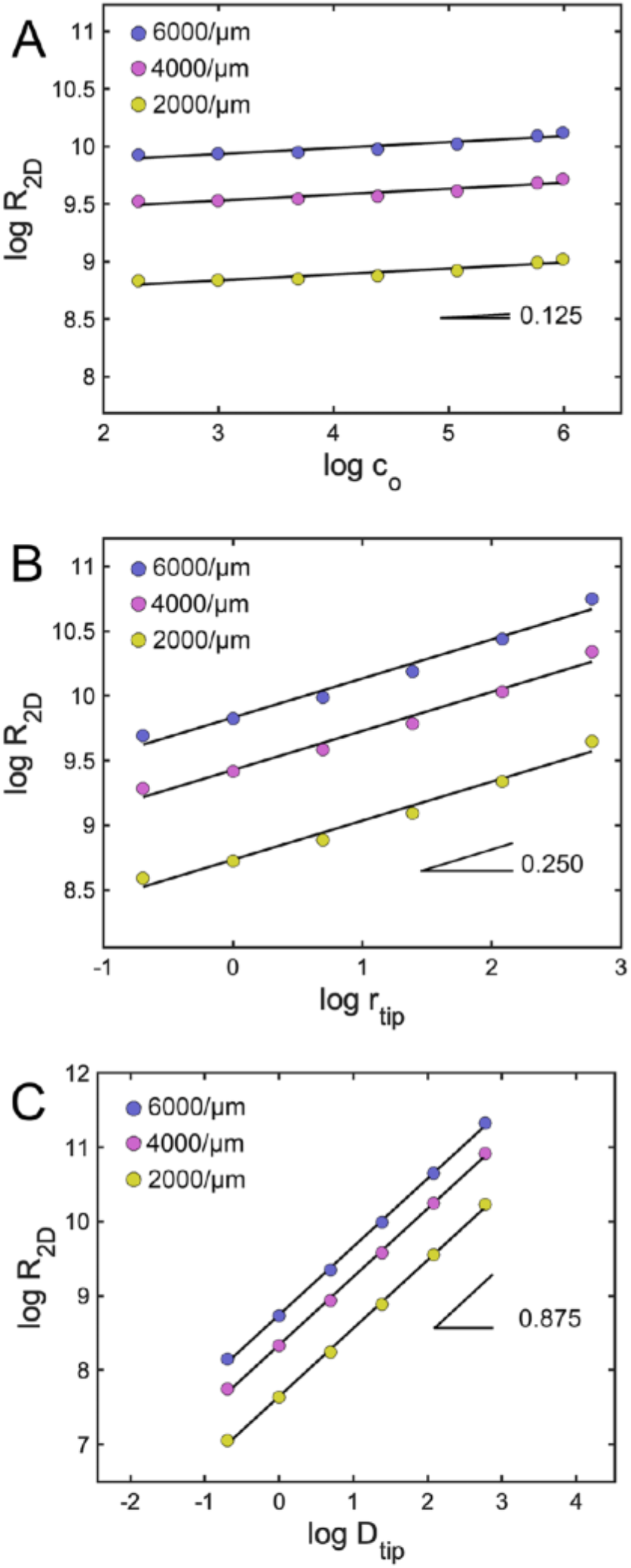
Log-log plots of 2D filament encounter rate (*R*_*2D*_) versus key parameter values. (A) *R*_*2D*_ vs. soluble profilin-actin concentration (*c*_*o*_). Slope: 0.125. (B) *R*_*2D*_ vs. filament capture radius (*r*_*tip*_). Slope: 0.25. (C) *R*_*2D*_ vs. tip diffusion coefficient (*D*_*tip*_). Slope: 0.875. Baseline parameter values: *c*_*o*_=100 µM; *r*_*tip*_=2 nm; *D*_*tip*_=4 µm^2^/sec.

## Notes

### Competing Interest Statement

The authors have declared no competing interest.

### Summary of Updates

We include two new approaches for calculating the rate of filament interaction with surface-bound monomers. Most of this new material is covered in Appendices C and D of the revise manuscript.

